# Limiting the loss of terrestrial ecosystems to safeguard nature for biodiversity and humanity

**DOI:** 10.1101/2021.02.07.428694

**Authors:** Jeremy S. Simmonds, Andres Felipe Suarez-Castro, April E. Reside, James E.M. Watson, James R. Allan, Pasquale Borrelli, Nigel Dudley, Stephen Edwards, Richard A. Fuller, Edward T. Game, Simon Linke, Sean L. Maxwell, Panos Panagos, Philippe Puydarrieux, Fabien Quétier, Rebecca K. Runting, Talitha Santini, Laura J. Sonter, Martine Maron

**Affiliations:** Centre for Biodiversity and Conservation Science, The University of Queensland, St Lucia 4072, Australia; School of Earth and Environmental Sciences, The University of Queensland, St Lucia 4072, Australia; Wildlife Conservation Society, Global Conservation Program, New York, United States of America; Institute for Biodiversity and Ecosystem Dynamics (IBED), University of Amsterdam, Amsterdam, The Netherlands; Department of Earth and Environmental Sciences, University of Pavia, Via Ferrata, 1, 27100 Pavia, Italy; Department of Biological Environment, Kangwon National University, Chuncheon 24341, Republic of Korea; Equilibrium Research, Bristol, United Kingdom; International Union for Conservation of Nature (IUCN), CH-1196 Gland, Switzerland; School of Biological Sciences, The University of Queensland, St Lucia 4072, Australia; The Nature Conservancy, South Brisbane 4101, Australia; Australian Rivers Institute, Griffith University, Nathan 4111, Australia; European Commission, Joint Research Centre (JRC), Ispra (VA), IT-21027, Italy; Biotope, 34140 Mèze, France; School of Geography, The University of Melbourne, Parkville 3010, Australia; School of Agriculture and Environment, The University of Western Australia, Crawley 6009, Australia

**Keywords:** Biodiversity, Convention on Biological Diversity, Framework Convention on Climate Change, Convention to Combat Desertification, ecosystems, post-2020, retention, Sustainable Development Goals, targets

## Abstract

Humanity is on a pathway of unsustainable loss of the natural systems upon which we, and all life, rely. To date, global efforts to achieve internationally-agreed goals to reduce carbon emissions, halt biodiversity loss, and retain essential ecosystem services, have been poorly integrated. However, these different goals all rely on preserving natural ecosystems. Here, we show how to unify these goals by empirically deriving spatially-explicit, quantitative area-based targets for the retention of natural terrestrial ecosystems. We found that at least 67 million km^2^ of Earth’s natural terrestrial ecosystems (~79% of the area remaining) require retention – via a combination of strict protection but more prominently through sustainably managed land use regimes complemented by restoration actions – to contribute to biodiversity, climate, soil and freshwater objectives under four United Nations’ Resolutions. This equates to retaining natural ecosystems across ~50% of the total terrestrial (excluding Antarctica) surface of Earth. Our results show where retention efforts could be focussed to contribute to multiple goals simultaneously. The retention targets concept that we present explicitly recognises that such management can and should co-occur alongside and be driven by the people who live in and rely on places where natural and semi-natural ecosystems remain on Earth.

## INTRODUCTION

Despite the dependence of humanity’s wellbeing on the natural world, we continue to erode nature, often irreversibly (IPBES 2019). While commitments to ‘sustainability’ abound, and have been at the core of international agreements since the seminal ‘Rio Declaration on Environment and Development’ almost 30 years ago (United Nations 1992), humans continue to degrade and destroy ecosystems at unsustainable rates in many parts of the world (Mackey et al. 2015; Watson et al. 2016; Thomas et al. 2017). This loss is accompanied by the finality of species extinction (Baisero et al. 2020), the erosion of ecosystem function and evolutionary processes (Watson et al. 2018), the loss of connection between people and nature (Ives et al. 2018), poorer-quality fresh water (Mapulanga & Naito 2019), loss of soil resources (Banwart 2011), depressed yields from living natural resources (Spera et al. 2020), and increased harm to people and nature from climate change (Maxwell et al. 2019). We lack a clear understanding of how much further biodiversity loss can occur before permanent, irrevocable damage is wrought on our life-support system (Maron et al. 2018a). However, evidence suggests we are at imminent risk of breaching (or have already exceeded) certain critical planetary thresholds (Steffen et al. 2015; Lenton et al. 2019). Given that the pressures on natural systems are accelerating as the human population grows and consumption intensifies (Leclère et al. 2020; Wiedmann et al. 2020), it is of urgent importance that we set a limit to the loss of natural ecosystems, to prevent further, irreparable damage.

The goals enshrined in multiple global agreements quite clearly depend upon retaining existing (and restored) natural and semi-natural ecosystems. For example, the global community’s aspirations regarding biodiversity conservation are comprehensively reflected in the stated objectives of the Convention on Biological Diversity (CBD), as well as various other ‘biodiversity-related conventions’ (e.g. CITES, Convention on the Conservation of Migratory Species of Wild Animals, World Heritage Convention, Convention on Wetlands (Ramsar)). Our reliance on, and need to conserve nature is further enshrined in the UN Sustainable Development Goals (e.g. Goal 15 “protect, restore and promote sustainable use of terrestrial ecosystems, sustainably manage forests, combat desertification, and halt and reverse land degradation and halt biodiversity loss”; also, Goals 6 (clean water), 14 (marine environment), and others). However, the targets underpinning the achievement of the goals captured in these agreements rarely articulate a desired, measurable outcome state–the amount of nature, where, and managed in what way, that is required for us to achieve particular biodiversity and sustainability objectives (Butchart et al. 2016; Elder & Olsen 2019).

Notably, dialogue relating to the proposed Post-2020 Global Biodiversity Framework under the CBD hints at an important shift, with an emphasis on *net* outcomes for biodiversity; namely, net improvements in ecosystems to 2050, delivered via an interim goal to increase the area, connectivity and integrity of natural ecosystems by at least 5% by 2030 (Secretariat of the Convention on Biological Diversity 2020). In large part, this goal’s achievement will necessitate limiting further losses of ecosystems–in other words, retaining the vast majority of what we have now. Some ongoing natural ecosystem depletion is inevitable to meet economic and social development imperatives (Zeng et al. 2020), underscoring the vital role of restoration activities to counterbalance losses using protocols such as offsets/ecological compensation (Simmonds et al. 2020). However, noting that restoration entails substantial time lags (Crouzeilles et al. 2016), is unfeasible or has uncertain outcomes in many instances (Gann et al. 2019), and that losses caused by pervasive threats (degradation by invasive species, illegal activities, etc.) rarely trigger compensation requirements (Maron et al. 2018b), a foundation of ecosystem retention is critical for achieving this ambitious outcome of a net increase in ecosystems extent (and net improvement in condition) by 2030.

International agendas that embed goals/targets which explicitly or implicitly require the retention of ecosystems have been largely implemented in isolation from one another, even with the emergence of the umbrella of the UN Sustainable Development Goals (CBD Subsidiary Body on Scientific Technical and Technological Advice 2019). This separation has predominated despite the fact that the requisite policies and on-ground actions needed to contribute to the achievement of the goals/targets captured among these conventions are likely to align in many instances (Secretariat of the Convention on Biological Diversity 2018; Jung et al. 2020; Soto-Navarro et al. 2020). The inefficiencies of this status quo - in terms of both missed opportunities for synergies, and the unification of agendas underpinned by a common response to managing nature including the retention of ecosystems – are now being recognised (CBD Subsidiary Body on Scientific Technical and Technological Advice 2019). Together with the opportunities presented by a move towards ambitious net outcomes-based goals in a post-2020 world (Bull et al. 2020; Secretariat of the Convention on Biological Diversity 2020), integrated delivery of various international agreements underscore the key role that the retention of ecosystems, framed by spatially-explicit ecosystem retention targets, could play in safeguarding our natural assets globally. Identification of retention targets – limits to loss – that set out what nature we need, and where, to achieve the full suite of goals associated with Earth’s remaining natural terrestrial ecosystems, has the potential to unite nature conservation and sustainable development goals across the international environmental agenda (Maron et al. 2018a).

Here, we present a method for quantifying ecosystem retention targets that are needed to achieve the nature-reliant ambitions of global policy agreements relating to biodiversity conservation, climate stabilisation, soil maintenance and water quality regulation. We set out an analytical approach for translating each of these goals to spatially-explicit guidance on how much and where existing natural terrestrial ecosystems should be retained to give us the best chance at meeting these various global objectives. We examine appropriate mechanisms for the management of these ecosystems, noting that a combination of strict protection and sustainable use will be required to align environmental, social and economic imperatives, rights and responsibilities. Further, we suggest that international cooperation will be crucial for determining country-level contributions to global ecosystem retention efforts.

## METHODS

### Global goals dependent upon natural terrestrial ecosystem retention

There are four current global agreements made under UN Resolutions for which natural terrestrial ecosystem retention is directly relevant: the Convention on Biological Diversity (CBD), the Framework Convention on Climate Change (UNFCCC), the Convention to Combat Desertification (UNCCD), and the Sustainable Development Goals (SDGs). These international agreements, each adopted and ratified by the vast majority of nations, contain clearly-articulated statements about what we expect from natural ecosystems, and are the logical starting point to mapping out how much terrestrial nature we need to retain, if we are to achieve each of these agreements’ goals.

We examined the goals, objectives, targets, and/or indicators under each of the four agreements to identify statements about desired outcomes that specifically depend upon (in whole or part) the retention of natural terrestrial ecosystems. This formed the basis of translating explicit quantifiable milestones and/or aspirational goals noted in the respective agreements into terrestrial ecosystem retention targets (Figure 1). Hereafter, we refer to the retention targets examined here that correspond to the four respective agreements as the ‘biodiversity conservation target’, ‘carbon storage target’, ‘soil maintenance target’ and ‘freshwater quality target’.

**Figure 1.**
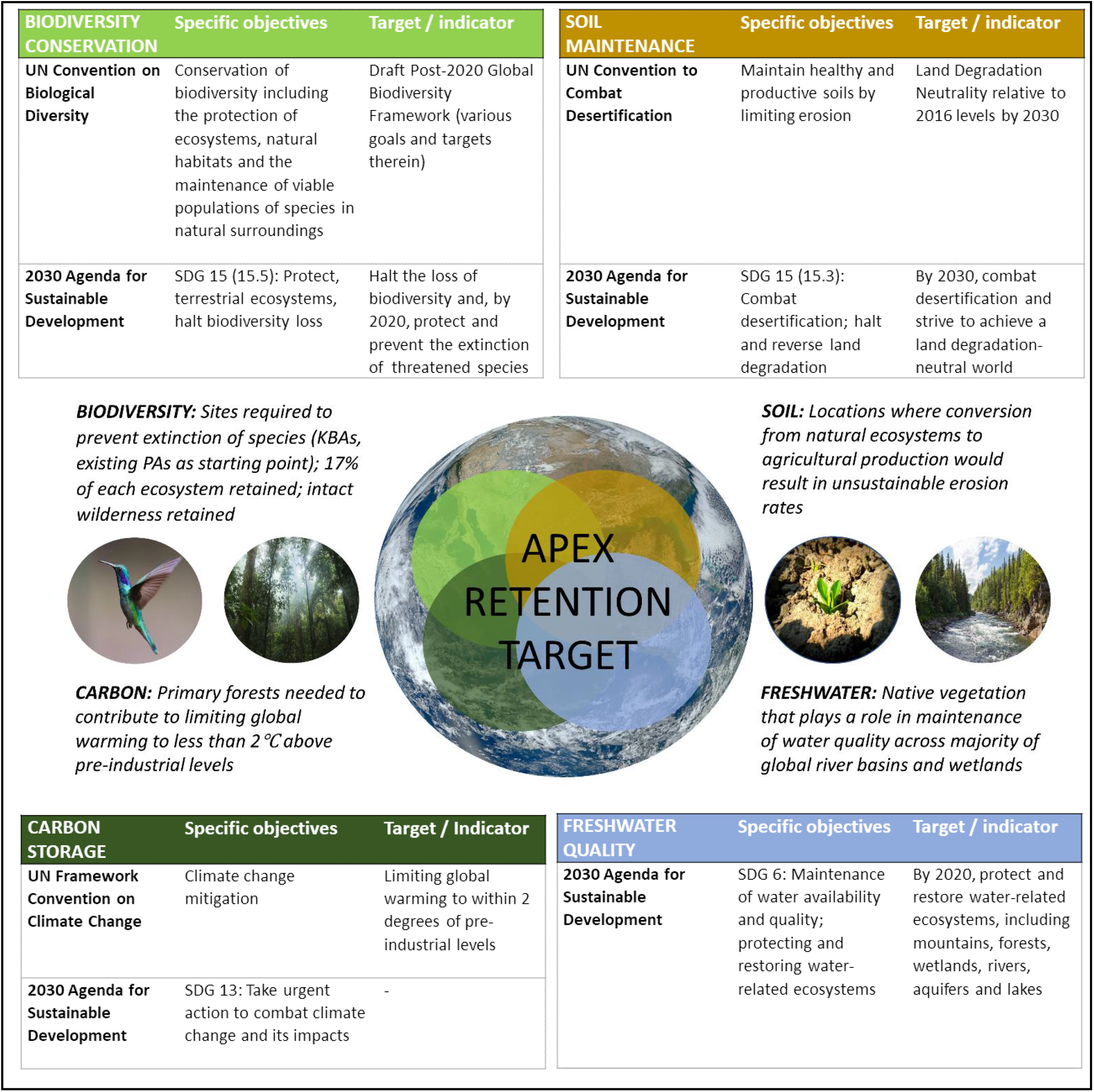
Objectives and associated targets/indicators that require retention of natural terrestrial vegetated ecosystems under international agreements. The basis for each agreement’s translation to the quantitative, spatially-explicit targets for the retention of natural terrestrial vegetated ecosystems that we examine, linked to the objectives/targets of each respective agreement, is outlined in the central part of the figure. A representation of the ‘apex retention target’ is also presented – this is made up of all the areas of natural terrestrial vegetated ecosystems that are captured by one or more of the four individual retention targets, and describes the amount of retention as a percentage of the total remaining terrestrial vegetated ecosystems on Earth (adapted from Maron et al. (2018a)).

In this analysis, we only considered areas for retention that are characterised by natural vegetation. While we refer here and throughout to ecosystems, we note that our maps select areas of natural land cover (albeit, along a spectrum of condition/human influence) categorised into broad classes, which are not of sufficiently high resolution to represent the full gamut of terrestrial ecosystems. We use the term ‘natural ecosystems’ throughout to underscore that this analysis, and the retention targets concept, is focussed on identifying and securing areas that are vegetated, and by proxy, support (or have potential to support) the suite of interacting and interconnected biotic and abiotic attributes (species assemblages and processes) that comprise ecosystems (notwithstanding issues of degradation – see Discussion). Importantly, the retention targets we propose here should be interpreted and considered in the context of ecosystem-specific targets, such as those being proposed for the Post-2020 Global Biodiversity Framework (Watson et al. 2020) and other complementary approaches to target-setting and planning (Dinerstein et al. 2020).

We did not include barren areas in our definition of natural ecosystems (e.g. ecosystems characterised by bare sand, exposed rock, ice and snow–including all of Antarctica). We acknowledge the value of these natural, largely non-vegetated environments, especially for biodiversity, but have excluded them here as this analysis is centred on one of the key actions needed to achieve multiple global goals – the retention of existing natural vegetation. Further, our analysis has a terrestrial focus, and does not extend to marine systems. Nonetheless, the framework we present for translating internationally-agreed goals to spatially-explicit quantitative ecosystem retention targets could be transferable to the marine realm. Amenable datasets may include marine ecosystems (Spalding et al. 2007; Spalding et al. 2012), human pressures (Halpern et al. 2015), carbon export in oceans (Henson et al. 2012; Roshan & DeVries 2017), and species ranges and protected area (various datasets produced by IUCN, BirdLife International, UNEP-WCMC; taxa-specific data (Kaschner et al. 2011)).

### Quantifying terrestrial ecosystem retention requirements

For each of the four retention targets – biodiversity conservation, carbon storage, soil maintenance, and freshwater quality – we produced maps that captured the amount and location of existing natural terrestrial vegetated ecosystems that need to be retained to contribute to the achievement of internationally-agreed goals. To represent natural ecosystems in this analysis (hereafter, we refer to this as the ‘natural ecosystems layer’), we used the MODIS Land Cover Type product MCD12Q1 at 250 m (native data resolution = 500 m) that was processed and modified by Borrelli et al. (2017). This layer, covering approximately 84% of Earth’s surface, is based on the International Geosphere Biosphere Programme (IGBP) system and reports seventeen land cover classes, including ten natural terrestrial vegetation classes (Supporting Information Table S1). We excluded non-vegetated and/or aquatic land cover types from our analysis (permanent wetlands, barren, snow/ice, water). Further, the three developed and mosaicked land classes (croplands, urban and built up, cropland/natural vegetation mosaics) were excluded from the analysis. For the ‘grasslands’ ecosystem category, we masked out grazing lands for the year 2000 using a spatial dataset that combines agricultural census data with satellite-derived land cover to map pasture extent (Ramankutty et al. 2008). The map we produced indicated that approximately 83.8 million km^2^ of natural terrestrial vegetated ecosystems remain on Earth (approximately 62% of the non-Antarctic land surface; noting that this does not account for the condition of this vegetation).

To determine how much and where natural terrestrial vegetated ecosystems need to be retained, we intersected existing mapping products linked to each of the four targets (described below) with the natural ecosystems layer. This allowed us to calculate the amount and location of natural terrestrial vegetated ecosystems that are spatially congruent with places recognised as having importance for each of the targets considered in this analysis. All spatial statistics were calculated using a Mollweide projection, with all mapping analyses being undertaken at a raster pixel resolution of approximately 250 m (i.e. at the resolution of natural ecosystems layer). An overview of the workflow for deducing spatially-explicit retention targets is presented in Figure 2.

**Figure 2.**
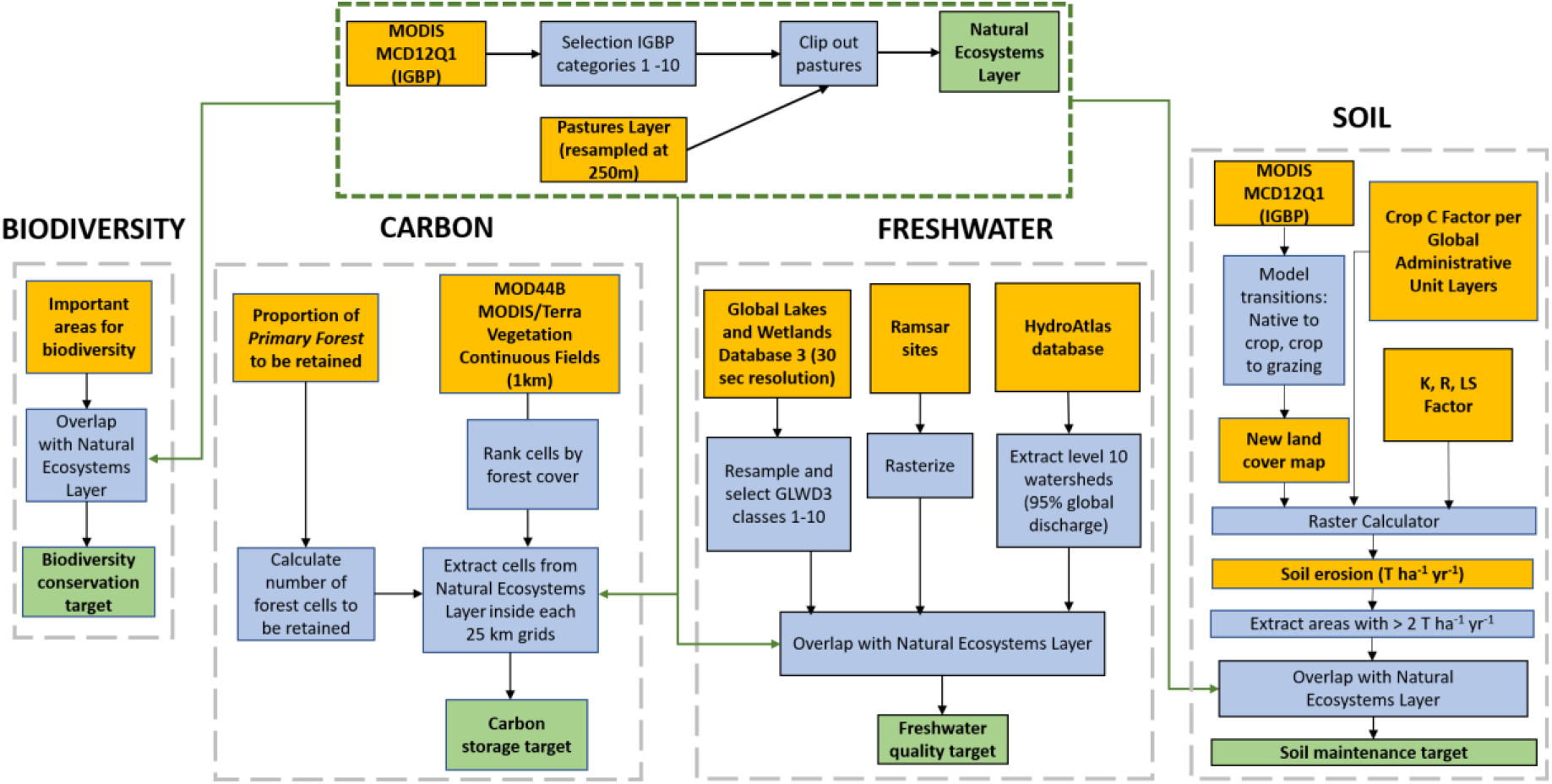
Overview of workflow for deriving maps for each of the four retention targets considered in this analysis. Yellow boxes represent input/derived datasets; Blue boxes represent GIS operations; Green boxes represent outputs. All raster inputs were resampled to match the resolution of the natural ecosystems layer.

#### Biodiversity conservation target

Language of the draft Post-2020 Global Biodiversity Framework under the CBD (August 2020), and the so-called Aichi Targets that it will supersede, is focussed on preventing extinctions, recovering threatened species, expanding the protection and management of important sites, and maintaining/enhancing ecosystem extent and resilience (including connectivity and intactness) (Secretariat of the Convention on Biological Diversity 2020). To translate these various aims to a biodiversity conservation retention target, we used the map produced by Allan et al. (2019), to identify natural terrestrial vegetated ecosystems that should be retained for multiple biodiversity conservation objectives (Figure 1). These maps assessed the spatial distribution and areal requirements needed to conserve 28,594 species, whilst also accounting for existing ecoregional representation targets and protected areas, Key Biodiversity Areas, and the maintenance of all large, contiguous areas with low human pressure (‘wilderness’). Detailed methods including the input data used to produce this map, and limitations on its interpretation, are provided Allan et al. (2019). We identified all natural terrestrial vegetated ecosystems from our natural ecosystems layer that are overlapped by this map, to identify the extent and distribution of natural ecosystems that require retention to contribute to the achievement of the biodiversity conservation objectives captured by the Allan et al. (2019) map. We assume that natural terrestrial vegetation retention in these places is integral to the persistence of biodiversity as represented by the Allan et al. (2019) analysis.

#### Carbon storage target

Forests play a key role in storing carbon. To determine how much and where natural ecosystems should be retained to stabilise levels of atmospheric carbon, we used spatial outputs from a harmonized land-use transition model for Shared Socioeconomic Pathway (SSP) 1 – a globally-accepted sustainable development pathway that respects environmental boundaries (van Vuuren et al. 2017). Spatial outputs for SSP1 are based on the IMAGE 3.0 integrated assessment model (IAM) (Stehfest et al. 2014) and describe annual transitions in different land-use categories (e.g. primary forest, secondary forest, pasture and cropland) between the years 1500 and 2100 at a spatial resolution of 0.5 x 0.5 degrees (approximately 25 km^2^ at the equator). The land use transitions capture subsequent effects on energy, water, and carbon exchanges between the land surface and the atmosphere, and are thus able to track the consequences of land use transitions on the global climate and carbon cycles. We chose SSP1 (of the five SSPs available (Riahi et al. 2017)) because the land-use transitions captured in this scenario have the greatest probability of limiting global warming to less than 2°C above pre-industrial levels (Hurtt et al. 2020).

We used spatial outputs for SSP1 to render a map of carbon retention through *primary forests* (defined as forested land that was not converted to an alternative land use type – urban, cropland, grazing land or secondary forest – between 1500 and 2050 according to the IMAGE 3.0 IAM (Hurtt et al. 2020); see below for explanation of focus on primary forest). The state of primary forest land in 2050 was subtracted from the state of primary forest land in 2015. The resulting index shows, for each 25 km^2^ grid cell, the proportion of primary forest required to be retained if forests are to effectively help stabilise levels of atmospheric carbon. We note that the index ascribes each 25 km^2^ grid cell a proportional retention value – that is, grid cells classed as 100% are not necessarily 100% covered by primary forest, but instead contain some primary forest in that cell, 100% of which must be retained to achieve the carbon storage target.

Once we identified the proportion of forested areas to be retained in each 25 km^2^ cell, we overlapped this with treed natural areas (forests, categories 1-5) from our natural ecosystems layer (Supporting Information Table S1). To allocate the spatial distribution of forest to be retained from this layer, we preferentially selected pixels starting with those with the highest coverage of trees based on the MOD44B Continuous Fields layer (an input into the MODIS-derived ecosystem layer used in this analysis). This is a 250 m spatial resolution biophysical parameter derived from the MODIS satellite. It reports annual estimates of the percentages of 1) surface vegetation cover, 2) bare soil, and 3) tree cover. The rationale for selecting forest pixels in this way was that pixels with a higher coverage of trees are more likely to be representative of primary forest; sparser areas on the other hand may be representative of more open (treed) ecosystem types or degraded forests. We continued this process until the area of forested pixels selected corresponded with the proportional retention value identified by the index of forest retention (to 2050) produced from SSP1 output at a 25 km^2^ grid cell resolution. To explore the implications of this for calculation of our ‘headline’ retention target, we repeated the sampling of forest pixels for the carbon goal preferentially selecting forest pixels that were also captured by at least one other of the biodiversity conservation, soil maintenance or freshwater quality target in this analysis. We found that the difference between the methods only considering the pixels with higher coverage of tree cover versus preferentially selecting forest pixels was less than 200,000 km^2^, which corresponds to less than 0.15% of the global area analysed.

Unlike the biodiversity, freshwater and soil goals in our analysis that span various natural terrestrial ecosystem types, our goal for carbon retention focuses only on primary forest ecosystems. This means that our representation of carbon retention will be an underestimate, particularly in parts of the world where other (non-forest ecosystems) play a key role in carbon storage. We note that the management (and retention) of other terrestrial ecosystem types (e.g. peatlands, mangroves, grasslands) can greatly benefit global efforts to mitigate climate change (Goldstein et al. 2020). However, we chose to focus on forested ecosystems as they contain the majority of terrestrial carbon stocks on Earth (Goldstein et al. 2020) and the spatial resolution of the study (approx. 25km^2^) precluded assessment of some carbon-dense ecosystems that have narrow or patchy geographic distributions (e.g. some mangrove or peatland ecosystems). Furthermore, we focused on primary forests as their effective management confers a multitude of co-benefits for other social and environmental values (Watson et al. 2018), and effectively managing carbon stocks in degraded forests (e.g. forests that are logged or used for livestock grazing) requires a complex set of management interventions that are beyond the scope of this study to capture.

The way we have identified forests for retention for the carbon goal should also be considered when interpreting the maps produced in this study – we reiterate that this study is not intended to be a high-resolution spatial prioritisation exercise to identify exactly where on the Earth’s surface native ecosystems need to be retained – rather, these maps are illustrative of the general location of where these ecosystems occur across the globe, noting that resolution mismatches of input layers prevented us from mapping forest retention for the carbon goal with the same degree of spatial precision as was the case for the other targets.

#### Soil maintenance target

The retention of natural ecosystems in areas where their removal would result in unsustainable rates of soil loss is crucial. Although several threats to land-based natural capital and soil quality are globally significant (salinization, acidification, nutrient depletion, contamination, waterlogging, etc.), for this analysis we focussed on soil erosion because it is a clear sign of land degradation that constitutes an irreplaceable loss and that cannot reasonably be restored. Indeed, soil erosion is the most immediate risk to land-based natural capital if natural vegetation is cleared.

Current global rates of soil loss by water erosion have been estimated by Borrelli et al. (2017), using the RUSLE-based (Revised Universal Soil Loss Equation) (Renard et al. 1997) modelling platform Global Soil Erosion Modelling (GloSEM). RUSLE incorporates rainfall erosivity and climate, erodibility of the soil, topography, and local farming systems and practices to predict the amount of soil lost (due to inter-rill and rill erosion processes) per unit area and time (t ha^-1^ yr^-1^) (Renard et al. 1997). Using data made available by Borrelli et al. (2017), we modelled the likely rate of soil loss in tons per ha per year if mapped natural terrestrial vegetated ecosystems were cleared. The possible new land use for 3,252 sub-national administrative units of 202 countries from the Global Administrative Unit Layers (United Nations Food and Agriculture Organization 2015), after clearing for agriculture, was either grazing or cropping depending on the main land use in the region according to Ramankutty et al. (2008). We assumed that administrative units with less than 50% coverage of grazing would be converted to the dominant current crop for that unit, whereas areas with more than 50% of coverage of grazing would be converted to grazed areas. The dominant crop per country was identified using data from the UN Food and Agriculture Organization (United Nations Food and Agriculture Organization 2016). Based on this land cover change, we then assigned values that measure the effect of cropping on the soil erosion process (C-Factors), based on thresholds identified by Borrelli et al. (2017). All grazing areas had the highest C-Factor values (0.5).

Sustainable or tolerable (Verheijen et al. 2009) soil erosion rates can be defined as rates of soil erosion that are equivalent to rates of soil formation consisting of mineral weathering as well as dust deposition. Soil formation rates vary widely across the globe depending on climate, geology, topography, and other factors (García-Ruiz et al. 2015). A global average soil formation rate of 2 t ha^-1^ yr^-1^ is an appropriate upper estimate of formation rates, based on calculation of rock weathering rates from exported water chemistry globally (Wakatsuki & Rasyidin 1992; Renard et al. 1997; Panagos et al. 2015). Using our map of areas where erosion rates were predicted to exceed 2 t ha^-1^ yr^-1^ if natural terrestrial vegetated ecosystems were replaced by agricultural land use, we identified areas from our natural ecosystems layer that should be retained to avoid unsustainable erosion, and thus contribute to land degradation neutrality goals.

#### Freshwater quality target

We produced a map of natural terrestrial vegetated ecosystems that should be retained to contribute to freshwater quality maintenance (e.g. via in-situ and catchment-level processes associated with filtration of impurities, reduction of sedimentation and pollutant run-off, etc.), to contribute to key global goals including SDG 6. To do this, we used three different datasets that map semi-aquatic (vegetated) ecosystems, areas of natural freshwater importance globally, and river basins that contribute disproportionately to global freshwater discharge. First, we used the Global Lakes and Wetlands Database 3 (GLWD) (30 Arc second resolution) (Lehner & Döll 2004). This dataset identifies areas that are predominantly water (e.g., lakes, rivers, marshes, flooded forests) and those that are partially water (e.g., intermittent lakes and wetland complexes). Here, we included classes 1-10 from the GLWD 3, but excluded classes 11 and 12 – these are attributed as covering less than 50% of the 30 Arc second raster grid upon which the GLWD 3 is based (Lehner & Döll 2004). To refine our analysis to natural terrestrial vegetated ecosystems associated with (overlapping) large semi-aquatic systems, we did not capture these small (<0.5 km^2^) ephemeral wetland complexes, noting that some such vegetation may have been captured in the complementary river analysis described below. We overlapped the layer produced using the GLWD 3 (classes 1-10) with our natural ecosystems map to identify terrestrial vegetated areas congruent with mapped wetlands. We undertook the same process for mapped (non-estuarine or marine) Ramsar sites, whereby natural ecosystems overlapped by Ramsar sites were selected.

To complement this ‘wetlands’ component of our freshwater quality target, we also used the HydroATLAS database developed by Linke et al. (2019) to identify river basins responsible for the majority of the planet’s freshwater discharge. HydroATLAS captures twelve nested levels of sub-basins at the global scale, each depicting consistently sized sub-basin polygons at scales ranging from millions (level 1) to tens of square kilometres (level 12). Using level 10 basins – a reasonable approximation of regional patterns of water discharge – we identified those basins collectively responsible for 95% of global freshwater discharge (approximately 75,000 individual basins). We then selected natural terrestrial vegetated ecosystems within these basins, under the rationale that these natural ecosystems are important for contributing to the regulation of water quality being discharged from these basins. To represent the amount and distribution of natural terrestrial vegetated ecosystems to be retained to contribute to global freshwater quality, we combined the areas of natural ecosystems selected for the GLWD overlay, Ramsar sites and river basin analyses (noting some overlap among the three).

### Mapping outputs and analysis

From these analyses, we produced four maps, displaying natural terrestrial vegetated ecosystems that should be retained to contribute to respective biodiversity conservation, carbon storage, soil maintenance and freshwater quality imperatives. On each map, we also showed existing natural ecosystems that were not captured by the respective retention targets. In addition to showing the distribution of natural ecosystem retention, these maps also allowed us to calculate the extent of natural ecosystems required to be retained to contribute to each of the four goals, as well as the percentage of natural ecosystems requiring retention compared to 1) the total extent of natural ecosystems remaining in 2012, and 2) the terrestrial surface of the planet (excluding Antarctica). We combined the four maps into a single global map to calculate an ‘apex retention target’ – in other words, the total amount (and percentage remaining) of natural terrestrial vegetated ecosystems that need to be retained to support the achievement of all four goals. To translate these findings to discrete units, we broke down natural ecosystem retention values by country.

## RESULTS

At least 67 million km^2^ (79%) of Earth’s remaining natural terrestrial vegetated ecosystems, covering approximately half the terrestrial (excluding Antarctica) surface of the planet, should be retained if internationally-agreed goals associated with biodiversity conservation, carbon storage, soil maintenance and freshwater quality are to be achieved (Table 1; Figure 3). This value represents the ‘apex retention target’ for natural terrestrial ecosystems – the amount needed for co-achievement of various environmentally-based goals. This target sets a limit to loss, and when combined with the spatial outputs from which it was derived, establishes (broadly) where we should aim to retain natural ecosystems, lest we risk compromising one or more of the environmental goals that nations of the world are committed to achieving.

**Figure 3.**
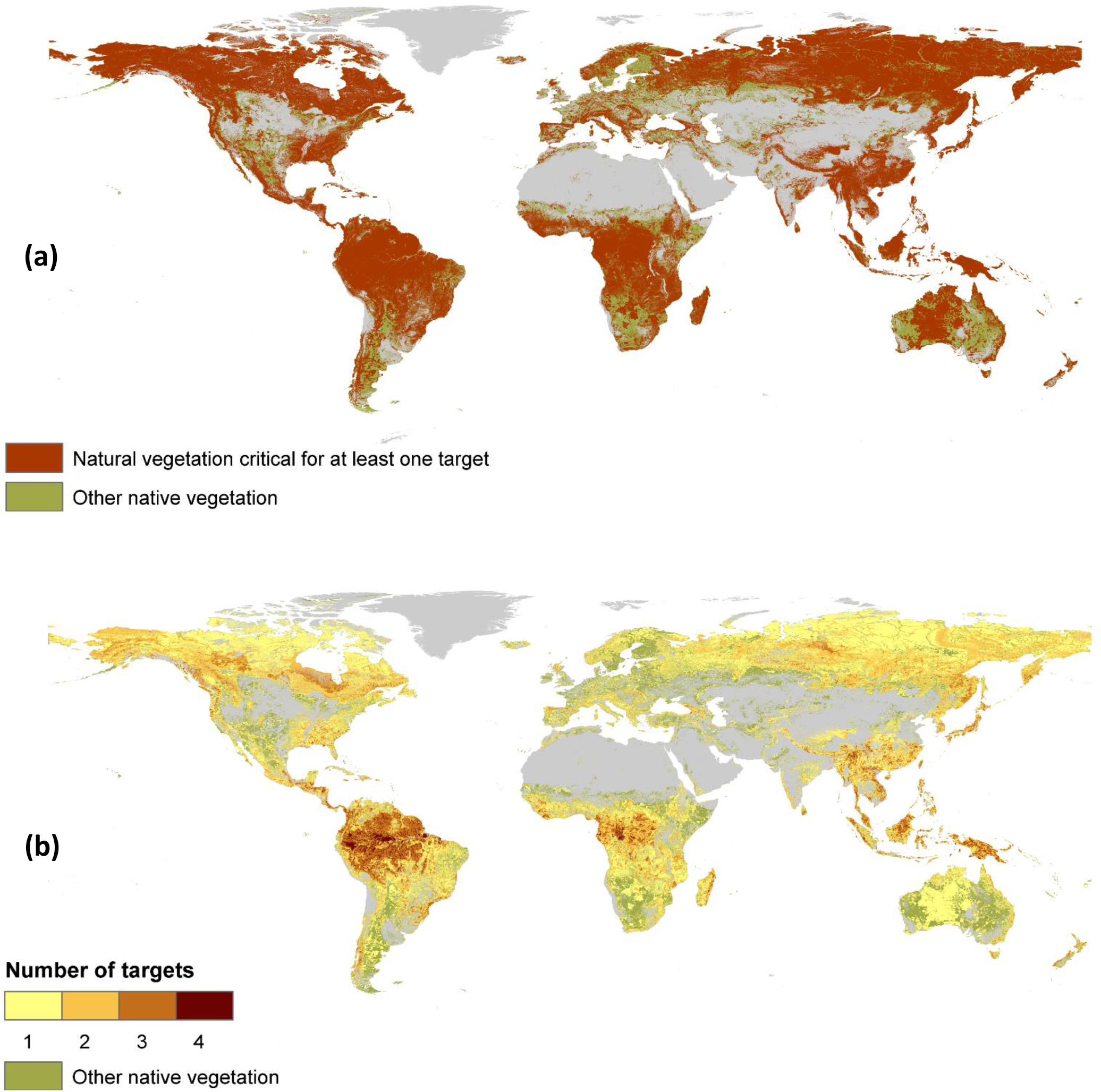
Natural terrestrial vegetated ecosystems requiring retention to contribute to the achievement of at least one target (a); and overlap of targets (b). Natural terrestrial vegetated ecosystems (‘native vegetation’) that is less critical for retention under any of the four targets are shown in light green. Grey shading represents areas of the land surface excluded from the analysis - either non-vegetated natural land cover types (e.g. rock, ice, barren), or where natural ecosystems have been replaced by other (anthropogenic) land cover types.

**Table 1.** Area of natural terrestrial vegetated ecosystems to be retained under each target (a); the area required to meet for all four targets (b). The percentage values do not add to 100%, as there are considerable areas of overlap of natural terrestrial vegetated ecosystems requiring retention.

| **a) Target** | **Area to be retained (km<sup>2</sup>)** | **Area as a % of remaining natural terrestrial vegetated ecosystems** | **Area as % of terrestrial Earth surface (excl. Antarctica)** |
| --- | --- | --- | --- |
| Biodiversity conservation | 43,551,955 | 51% | 32% |
| Carbon storage | 20,984,782 | 25% | 15% |
| Soil maintenance | 36,493,119 | 43% | 27% |
| Freshwater quality | 12,133, 826 | 14% | 9% |
| **b) Overlap among targets** |  |  |  |
| Number of targets contributed to |  |  |  |
| 1 | 34,430,437 | 41% | 26% |
| 2 | 22,314,968 | 26% | 17% |
| 3 | 9,263,759 | 11% | 7% |
| 4 | 1,499,633 | 2% | 1% |
| **Total** | **67,508,798** | **79%** | **50%** |

Biodiversity conservation required the largest extent of natural ecosystem retention (Table 1; Figure 4), with over 43 million km^2^ (51% of remaining ecosystems) required to contribute to this goal with its various objectives (species persistence, ecosystem representation, securing important sites such as existing KBAs, retention of contiguous areas of low human pressure (‘wilderness’)). There was relatively low overlap among the spatial distribution of retained ecosystems satisfying multiple goals – approximately 27% of retained ecosystems contributed to two targets, while only 2% contributed to all four targets (Table 1).

**Figure 4.**
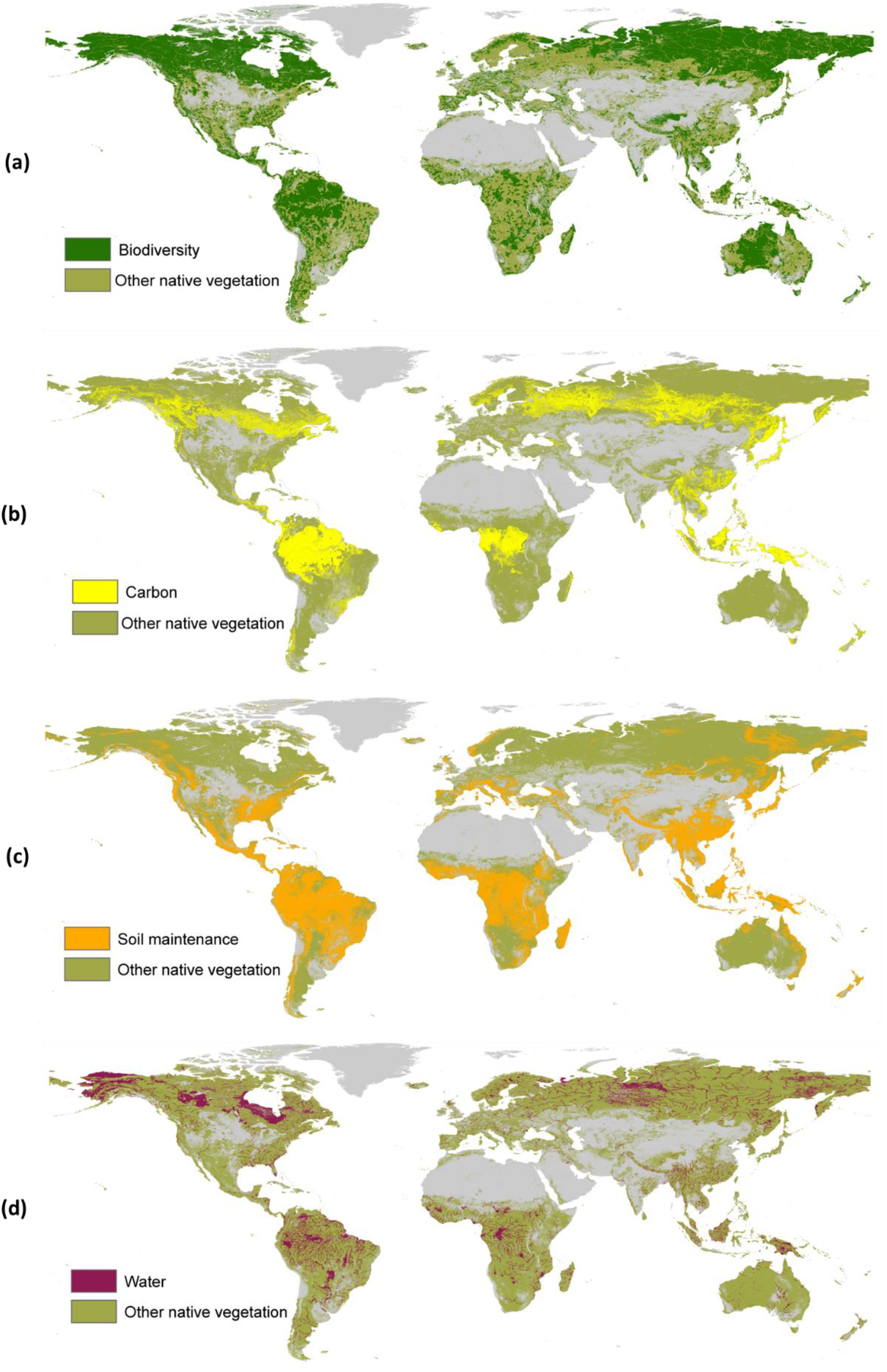
Natural terrestrial vegetated ecosystems required for retention to meet each of the four targets: biodiversity conservation (a); carbon storage (primary forest only) (b); soil maintenance (c); and freshwater quality (d). Natural terrestrial vegetated ecosystems (‘other native vegetation’) is shown in light green.

Individual countries differed markedly in their ecosystem retention requirements (Figure 4a-d; Figure 5). The area for retention required to meet the apex target of 79% is disproportionately shared among several large countries, with Russia, Canada, Brazil, United States and Australia accounting for more than half (52%) the extent alone. Of the 66 countries with ecosystem retention targets of at least 90% of existing natural terrestrial ecosystem extent, the vast majority (n=63) are in Africa, Asia and the Americas. Imperatives to meet national socio-economic goals are likely to be particularly acute in many such countries, presenting a conflict between the achievement of global environmental goals, and development.

**Figure 5.**
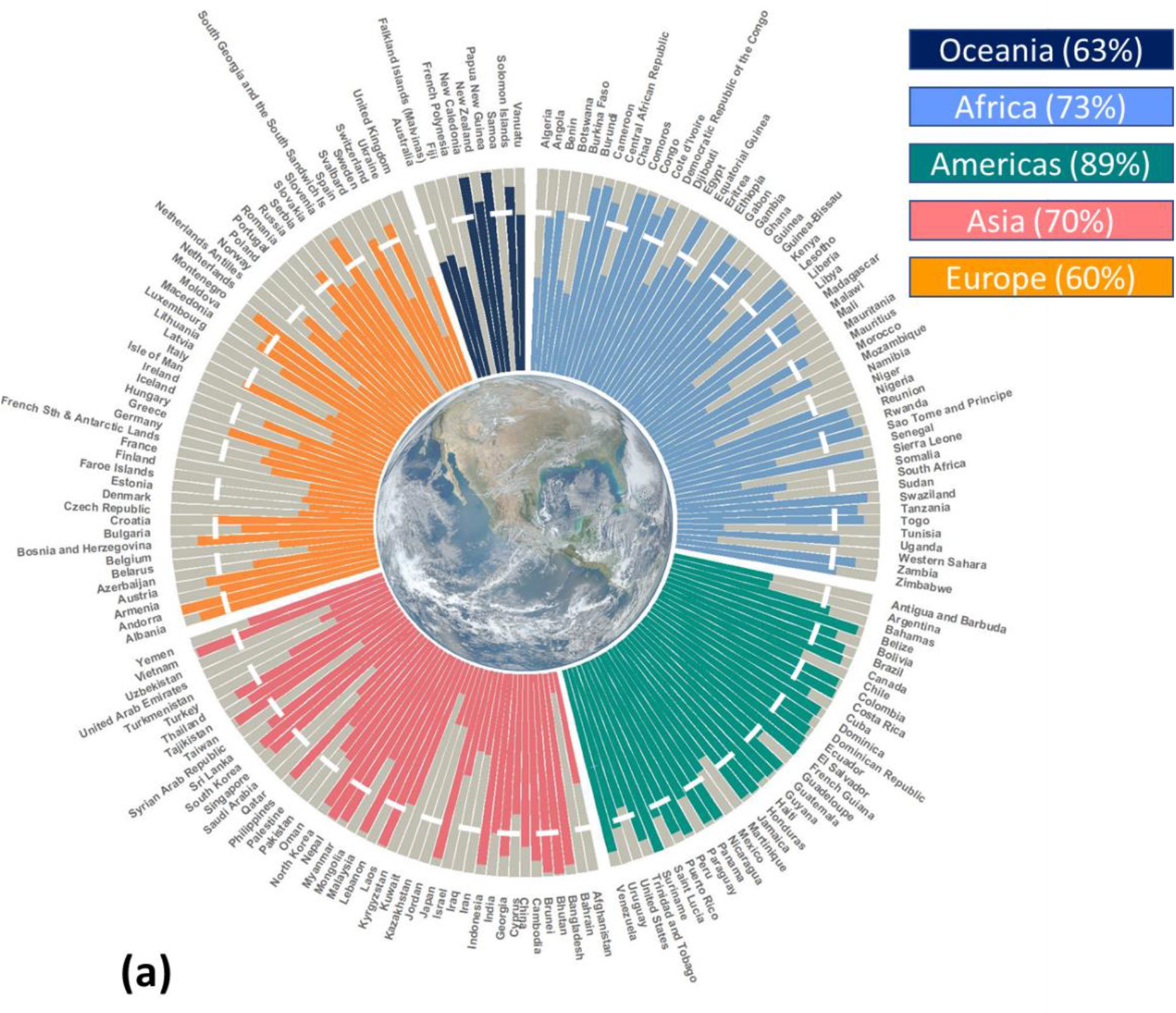
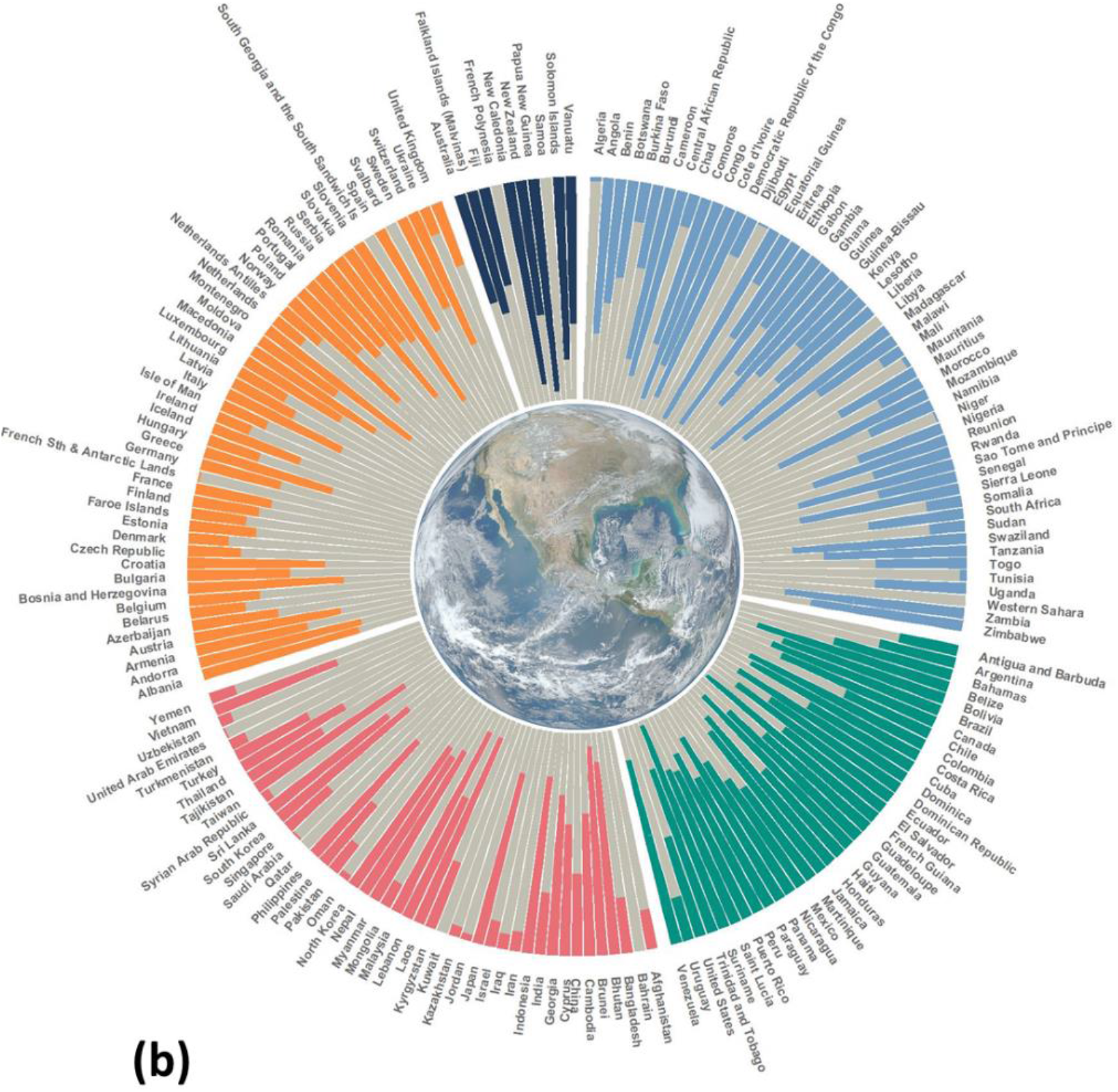
Country-level ecosystem retention values (coloured section of bars) as proportion of total amount of natural terrestrial ecosystems remaining. The white dashed line represents the apex global retention target of 79%. Percentage values in the key are the average natural terrestrial ecosystem retention value per continent (a). Amount of natural terrestrial ecosystems requiring retention as a percentage of total land area of each country (b). Countries with land area greater than 500 km^2^ shown here for display purposes, with all national retention values presented in Supporting Information Table S2. Overseas dependencies are classified according to their country (and continent) of association.

## DISCUSSION

Our analysis indicates that at least 79% of the remaining extent of natural ecosystems should be retained, with any loss from the areas we identify for retention potentially compromising our ability to achieve globally-agreed environmental goals of humanity. This equates to keeping at least half of the planet under natural vegetation coverage - a striking finding that aligns with other recent analyses that call for greatly increased ambition to conserve at least half of the natural world (e.g. Dinerstein et al. 2020; Wilson 2016). Importantly, achieving such an ambitious outcome (a global retention target of 79%) is not predicated on the exclusion of people from natural and semi-natural landscapes, nor should it compromise development imperatives - indeed, changes in the way we manage and draw from the biosphere can see this ambition become a reality (Tallis et al. 2018). This retention-centred approach to framing and establishing global environmental targets that we present sets a limit to the loss of nature. This is vital in a world where the depletion of natural systems is rampant - for example, over 4 million km^2^ of land was converted to human land uses between 1993 and 2009 (Watson et al. 2016), while up to a million species are threatened with extinction (IPBES 2019).

### Unifying multiple goals

The retention targets approach we demonstrate can help achieve multiple goals under a unified approach - so far, an elusive prospect (Scharlemann et al. 2020). It addresses key points of criticism of previous global targets, such as the Aichi Targets (and especially Target 11) under the CBD with their lack of focus on outcomes (Barnes et al. 2018). Moreover, it maps a path towards integrating conservation with the various expectations we have of nature (e.g. service provision), which has been lacking from global endeavours such as those encapsulated in the UN Sustainable Development Goals (Zeng et al. 2020). This is because it points to a scientifically-formulated, spatially-explicit and measurable outcome state, which is aligned with the goals of these various agreements.

Our method, founded on the notion of setting a limit to loss via retention targets, offers a complementary approach to various other proposals for setting global environmental targets, in that it establishes what we need (the desired outcome) which can then lay a foundation for how to get there (mechanisms). One such example is the ‘Global Safety Net’, which provides a spatially-explicit representation of where vastly increased conservation efforts need to be concentrated, to achieve complementary biodiversity and carbon objectives (Dinerstein et al. 2020). Based upon a detailed suite of biodiversity and carbon data, the outputs of this analysis provide a scientifically-robust roadmap for future land use planning and decision-making. We concur with Dinerstein et al. (2020) that protected areas will always be the key cornerstone of conservation efforts, but that they alone cannot preserve species, ecosystems and ecological functions (unless the ambition for protected area coverage is dramatically and perhaps implausibly increased). Indeed, our results demonstrate that we need vastly more nature retained than is contained within current protected areas, many of which are only ‘paper parks’ (Di Minin & Toivonen 2015). Even ambitions to increase their coverage will, in the absence of a more holistic framing, potentially fall short of securing all of the ecosystems we must keep for the full gamut of services that nature provides us, as well as providing sufficient space for wild species to thrive, and for evolutionary processes to continue. This underscores the necessity of framing environmental targets in the retention (and restoration of) natural systems across tenures and land use regimes.

Interestingly, our results revealed limited overlap between multiple targets in some parts of the world - a likely function of our input datasets being constrained to certain elements of the biota (e.g. ecoregional representation targets for biodiversity, primary forest only for carbon, focus of vegetation retention in subset of river basins). Given that our approach is a conservative protocol for identifying sites for retention, we stress that opportunities for synergies must be carefully considered. Our analysis is not an optimisation, and differs from spatial prioritisations. Jung et al. (2020) used an optimisation approach to identify high priority sites for species conservation, carbon retention and water provisioning. This approach, which identified priority sites but did not establish the amount and spatial distribution of land needed to meet specific outcomes-based targets, is another valuable complement to our retention target maps - especially when it comes to identifying what and where particular actions should be most urgently undertaken. For example, natural ecosystems identified by our study as warranting retention, and by Jung et al. (2020) to be of high priority, might warrant more targeted interventions (strict protection, carefully managed use to allow for provisioning of and access to ecosystem services). Conversely, lower priority sites may be more amenable to sustainably-managed mixed use (e.g. intensification of existing agricultural practice) landscapes, albeit, where the focus is on long-term ecosystem retention (complemented by restoration, which will be especially critical for depleted habitats for many threatened species).

### Implementation - opportunities and considerations

As noted elsewhere (Maron et al. 2018a; Dinerstein et al. 2020), achieving retention targets will require a carefully developed mixture of protection and sustainable management of natural and semi-natural vegetation (in parallel with restoration of degraded sites) beyond strictly protected areas, alongside transformative changes in consumption patterns (supply and demand of food and resources) (Leclère et al. 2020). Many pragmatic decisions and some trade-offs will be needed. Well-managed protected areas are long-standing mechanisms for securing ecosystems (Maxwell et al. 2020), which can be complemented by newer approaches such as other effective area-based conservation measures (OECMs) (Dudley et al. 2018), including privately held land (Clements et al. 2018). The stewardship of Indigenous people and other local communities with connection to the land and knowledge of its management, will be necessary for the preservation of large swathes of natural and semi-natural areas, where such retention is aligned with the aspirations of Indigenous custodians (Garnett et al. 2018; O’Bryan et al. 2020). Security of tenure and access is an essential enabling condition for this stewardship to be equitable and undertaken in a manner that is consistent with local peoples’ aspirations. Strategies favouring commercial land uses that retain important natural/semi-natural vegetation offer a further pathway to retention; for example, sustainable forest management (e.g. via certification schemes (Kleinschroth et al. 2019)) or wildlife-sympathetic and nomadic livestock grazing (Fynn et al. 2016; Liao et al. 2020), as compared with conversion to intensive single-crop agriculture.

Other effective levers, such as moratoria on conversion of sensitive ecosystems, are increasingly applicable to commercial products and their supply chains (Newton et al. 2013). Jurisdictional law and policies regarding the use and management of natural resources also have a central role to play in framing where and to what extent certain elements of the biota can be utilised (e.g. harvest limits (Di Minin et al. 2019)), and in imposing environmental conditions on permits for certain activities, either through sectorial policies and regulations, or following an environmental impact assessment (EIA) process, which is well-established globally. One of EIAs central tenets, the mitigation hierarchy, is a globally-ubiquitous instrument legislated by governments, and required by many financiers and industries, to minimise the impacts of development and balance environmental harm with equivalent gains elsewhere (zu Ermgassen et al. 2020). Harnessing the corporate sector generally (e.g. through commitments to ‘science-based targets’ (Andersen et al. 2020)), and the mitigation hierarchy specifically (Simmonds et al. 2020) holds substantial promise for resolving tensions between development and ecosystem conservation.

The retention target framing explicitly recognises that much of the nature we need is where we live, work, produce our food and extract resources, and that keeping this nature *in situ* necessitates a multi-faceted approach. However, we note that for any given site for which retention of natural ecosystems is required, the management and governance standards needed will differ depending on the objective to which that place contributes. Where ecosystem retention contributes to multiple goals, the strictest standard required to ensure adequate management should apply. For example, a site for which retention contributes to both biodiversity and carbon goals might be amenable to sustainable production (e.g., grazing of livestock) from a carbon perspective, but this production may unsustainably affect its biodiversity values. Here, targeted management to maintain the biodiversity values would be necessary, extending to strict protection if other land uses were incompatible with conserving the site’s biodiversity values.

A pitfall of global environmental targets is that they tend to be taken *prima-facie* without a clear understanding of the assumptions and limitations inherent to the target-setting, especially as this applies to their implementation on the ground (Garcia et al. 2020). Ultimately, achieving targets such as the retention concept we present, requires changes in behaviours and decision-making at multiple scales by many different people. In fact, the way we make decisions on the targets, and the role of human agency in this process, is likely more important than the targets themselves (Garcia et al. 2020). In the context of retention targets, this particularly matters as countries (and communities therein, especially Indigenous communities) that support a high proportional extent of ecosystems requiring retention will carry a disproportionate burden when it comes to achieving a global apex target such as the one we present in this analysis. Maron et al. (2020) suggested that the loss of nature in many such countries will profoundly affect local people where there may be a more direct reliance on ecosystem services (clean water access, for example), in the absence of extensive and maintained infrastructure. Despite the local (and broader regional and global benefits) of keeping nature *in situ*, an expectation that such peoples and nations retain the vast majority of their natural ecosystems presents complex challenges, due to issues around power imbalances, governance, capacity and competing (development) aspirations (Maron et al. 2020). Recognising that broad-scale, intensive development is but one of several key pathways to improved socio-economic conditions, the support (finance, capacity and empowerment; distribution of the cost of retention) of other (external) actors may be needed in some instances (but only if requested) to ensure that development can occur sustainably in these places, while not compromising local ecosystem service provision nor global environmental goals (Maron et al. 2020). Meaningful engagement with rights-holders and other stakeholders at multiple scales, from the outset of any decision-making on development and retention objectives, will be essential to these endeavours.

Establishing an apex environmental target for global efforts to be directed towards does have limitations. For example, Purvis (2020) contended that a single apex target focussed on biodiversity is unwise, given that the different motivations for conserving biodiversity (e.g., preserving intrinsic vs utilitarian values) need fundamentally different strategies. We acknowledge this, but also recognize that the global agreements on which our analysis is based provide the most holistic and pluralistic set of goals for scientifically-formulated targets for the retention of natural ecosystems. In this respect, our apex target is relevant to the policy processes that multilateral environmental agreements trigger among its signatories. For example, such a target would act as a useful empirical marker for the achievement of overarching aspirations of these agreements, such as the CBD’s 2050 Vision of humanity “living in harmony with nature”.

### Limits to loss

Our analysis acts as a starting point for establishing where and what nature should be retained to help us meet global environmental commitments. We make a range of assumptions - most notably, that all selected areas of natural vegetation are of equal value (and condition) with respect to their contribution to the goals they are selected under. Variability in condition (and threats) has substantial consequences for how sites are managed and retained - in the case of biodiversity for example, extensive pressure-free lands will have different management imperatives compared with habitat remnants in human mosaics. Improving the inputs for selecting sites for retention, and the resolution of the ecosystem data (including information on condition (e.g., Hansen et al. 2020), may allow for a more refined retention map and target value to be produced. This, however, may be more suited to regional-or local-level analyses, where higher resolution data inputs are available, and where relevant local actors can be involved in setting targets and deciding how to achieve them. We reiterate that this analysis sets a minimum threshold for retention, given that we did not comprehensively account for all elements of the biota linked to each goal (e.g., our carbon target only deals with primary forest; we did not consider biodiversity in arid (barren) environments), as well as the relatively narrow scope of goals that we considered; additional goals can only increase this value.

With ongoing losses a certainty, and numerous questions around the capacity of many ecosystems to recover (at least in timeframes congruent with human lifetimes/several generations, let alone time-bound international environmental agreements), balancing losses with gains to achieve net outcomes cannot be relied upon to maintain ecosystems at the required retention amount. Moreover, the extensive degradation within remaining natural ecosystems - almost 60% of forests globally have compromised ecological integrity and are in a degraded state (Grantham et al. 2020) - further underscores that the retention values we present should be considered as conservative targets. Retaining vegetation well above the apex target will act as a buffer for the (inevitable) net losses incurred as a result of continued land use change, resource extraction and degradation. Of course, a small amount of net loss could be absorbed in areas that our maps indicate are less critical for retention for any of the four targets we consider (~21% of remaining). However, we urge great caution in this interpretation, as the value of these sites for other environmental and/or sustainable development imperatives, as well as the ecosystem services they may supply to proximal communities, has not been assessed here. Thus, we strongly advise that these ecosystems should not be considered ‘worthless’ regarding their contribution to nature and people, with any actions in them (i.e. development) subject to rigorous scrutiny. This especially applies where the loss of such ecosystems will negatively affect the supply of services to local communities, albeit without necessarily detracting from the achievement of goals that are far broader in their scope.

Increasingly, nations and other influential actors are coalescing around the idea that transformative change to the way in which we live, and manage our biosphere, is needed. Noting that time is running out to halt and reverse the degradation of nature, we propose that a good starting point for initiating this change is asking the simple question: what do we need and want from nature? This question manifests here in the form of a retention target for a least four of nature’s contributions to people, which can allow us to map the places and inform the actions needed to maintain natural ecosystems. For the benefit of biodiversity, and for people too, we need to keep a great deal of the world’s remaining natural ecosystems in place. This analysis provides a starting point to the questions of ‘where’ and ‘how much’, this should be.

## ACKNOWLEDGEMENTS

Pasquale Borrelli is funded by the EcoSSSoil Project, Korea Environmental Industry & Technology Institute (KEITI), Korea (Grant No. 2019002820004).

## AUTHOR CONTRIBUTIONS

The data and methods for the spatial quantification of retention targets presented in this paper were prepared and developed by MM, JSS, JEMW, AFSC, AER, JAA, SLM, PB, PPa, SL, TS, LJS and RKR. Spatial analyses were conducted by AFSC and AER. JSS and MM wrote the manuscript. All authors contributed to the conceptualisation of this paper, to subsequent drafting and editing of the paper, and approved its final version.

