## Supporting Information for "Limiting the loss of terrestrial ecosystems to safeguard nature for biodiversity and humanity"

**Supporting Information Table S1**

**Supporting Information Table S2**

**Table S1. Land cover classes defined by the International Geosphere Biosphere Programme (IGBP), which includes 10 natural terrestrial ecosystem classes (Indices 1 -10), three developed and mosaicked land classes, and four non-vegetated land classes. Only categories 1 -10 were considered to calculate the total amount of natural ecosystems to be retained in this analysis.**

| Index | Surface Type | Description |
|---|---|---|
| 1 | Evergreen Needleleaf Forests | Lands dominated by trees with a percent canopy cover >60% and height >2 m |
| 2 | Evergreen Broadleaf Forests | Lands dominated by trees with a percent canopy cover >60% and height >2 m |
| 3 | Deciduous Needleleaf Forests | Lands dominated by trees with a percent canopy cover >60% and height >2 m |
| 4 | Deciduous Broadleaf Forests | Lands dominated by trees with a percent canopy cover >60% and height >2 m |
| 5 | Mixed Forests | Lands dominated by trees with a percent canopy cover >60% and height >2 m. Consists of tree communities with interspersed mixtures or mosaics of the other four forest cover types. None of the forest types exceeds 60% of landscape. |
| 6 | Closed Shrublands | Lands with woody vegetation <2 m tall and with shrub canopy cover >60%. The shrub foliage can be either evergreen or deciduous. |
| 7 | Open Shrublands | Lands with woody vegetation <2 m tall and with shrub canopy cover between 10%-60%. The shrub foliage can be either evergreen or deciduous. |
| 8 | Woody Savannas | Lands with herbaceous and other understory systems, and with forest canopy cover between 30%-60%. The forest cover height >2 m. |
| 9 | Savannas | Lands with herbaceous and other understory systems, and with forest canopy cover between 10%-30%. The forest cover height >2 m. |
| 10 | Grasslands | Lands with herbaceous types of cover. Tree and shrub cover is <10%. |
| 11 | Permanent Wetlands | Lands with a permanent mixture of water and herbaceous or woody vegetation that cover extensive areas. The vegetation can be present in either salt, brackish, or fresh water. |
| 12 | Croplands | Lands covered with temporary crops followed by harvest and a bare soil period (e.g., single and multiple cropping systems.)<br>Note that perennial woody crops will be classified as the appropriate forest or shrub land cover type. |
| 13 | Urban and Built-Up | Land covered by buildings and other man-made structures. |
| 14 | Cropland/Natural Vegetation Mosaics | Lands with a mosaic of croplands, forest, shrublands, and grasslands in which no one component comprises more than 60% of the landscape. |
| 15 | Snow and Ice | Lands under snow and/or ice cover throughout the year. |
| 16 | Barren | Lands made up of exposed soil, sand, rocks, or snow which never have >10% vegetated cover during any time of the year. |
| 17 | Water | |

**Table S2. Amount of natural ecosystem retention as a percentage of total remaining natural ecosystems, and land area, for 232 nations and overseas dependencies. These results should be interpreted with care (especially for small nations/dependenices), noting the coarse resolution of the input date underpinning this global-level analysis.**

| Country | Area ($km^2$) | Total natural ecosystems ($km^2$) | Retention (% remaining natural ecosystems) | Retention (% land area) |
|---|---|---|---|---|
| Afghanistan | 643557 | 203167 | 56% | 18% |
| Albania | 28681 | 22594 | 92% | 73% |
| Algeria | 2326150 | 123372 | 46% | 2% |
| American Samoa | 165 | 85 | 0% | 0% |
| Andorra | 507 | 380 | 100% | 75% |
| Angola | 1254950 | 1154679 | 76% | 70% |
| Anguilla | 92 | 22 | 64% | 15% |
| Antigua and Barbuda | 540 | 291 | 50% | 27% |
| Argentina | 2787410 | 2001696 | 49% | 35% |
| Armenia | 29703 | 19157 | 79% | 51% |
| Aruba | 201 | 67 | 64% | 21% |
| Australia | 7718920 | 6626812 | 60% | 51% |
| Austria | 83896 | 65760 | 85% | 67% |
| Azerbaijan | 86037 | 55464 | 61% | 39% |
| Bahamas | 12192 | 8198 | 81% | 55% |
| Bahrain | 643 | 26 | 10% | 0% |
| Bangladesh | 138505 | 35716 | 95% | 25% |

| **Country** | **Area (km²)** | **Total natural ecosystems (km²)** | **Retention (% remaining natural ecosystems)** | **Retention (% land area)** |
|---|---|---|---|---|
| Barbados | 449 | 276 | 43% | 26% |
| Belarus | 207316 | 153086 | 33% | 25% |
| Belgium | 30612 | 20957 | 47% | 32% |
| Belize | 22149 | 20462 | 97% | 90% |
| Benin | 116922 | 81846 | 80% | 56% |
| Bermuda | 20 | 6 | 53% | 15% |
| Bhutan | 39991 | 34982 | 99% | 87% |
| Bolivia | 1092610 | 958112 | 88% | 77% |
| Bosnia and Herzegovina | 51538 | 41790 | 87% | 70% |
| Botswana | 581127 | 549266 | 46% | 44% |
| Brazil | 8523590 | 7341699 | 94% | 81% |
| British Virgin Islands | 116 | 55 | 59% | 28% |
| Brunei | 5785 | 5533 | 98% | 94% |
| Bulgaria | 111083 | 77846 | 65% | 46% |
| Burkina Faso | 274009 | 121193 | 40% | 18% |
| Burundi | 27368 | 11156 | 94% | 38% |
| Cambodia | 182847 | 127310 | 92% | 64% |
| Cameroon | 467816 | 385598 | 96% | 79% |
| Canada | 9923600 | 7910836 | 95% | 75% |

| Country | Area (km²) | Total natural ecosystems (km²) | Retention (% remaining natural ecosystems) | Retention (% land area) |
|---|---|---|---|---|
| Cayman Islands | 210 | 103 | 87% | 43% |
| Central African Republic | 622659 | 600145 | 89% | 86% |
| Chad | 1279120 | 361506 | 53% | 15% |
| Chile | 745768 | 414651 | 76% | 42% |
| China | 9388280 | 4055541 | 85% | 37% |
| Christmas Island | 125 | 6 | 48% | 2% |
| Cocos (Keeling) Islands | 18 | 1 | 43% | 2% |
| Colombia | 1142650 | 1032600 | 98% | 88% |
| Comoros | 1727 | 1223 | 97% | 69% |
| Congo | 346334 | 333542 | 98% | 94% |
| Cook Islands | 151 | 92 | 0% | 0% |
| Costa Rica | 51375 | 44721 | 99% | 86% |
| Cote d'Ivoire | 323412 | 276507 | 90% | 77% |
| Croatia | 55889 | 43856 | 79% | 62% |
| Cuba | 109727 | 66846 | 88% | 54% |
| Cyprus | 9158 | 7872 | 69% | 59% |
| Czech Republic | 78668 | 50361 | 37% | 24% |
| Democratic Republic of the Congo | 2342080 | 2209656 | 96% | 91% |

| Country | Area ($km^2$) | Total natural ecosystems ($km^2$) | Retention (% remaining natural ecosystems) | Retention (% land area) |
|---|---|---|---|---|
| Denmark | 42605 | 23882 | 32% | 18% |
| Djibouti | 21568 | 7094 | 57% | 19% |
| Dominica | 769 | 704 | 97% | 89% |
| Dominican Republic | 48626 | 35322 | 96% | 70% |
| Ecuador | 257026 | 224281 | 98% | 86% |
| Egypt | 1002450 | 9294 | 69% | 1% |
| El Salvador | 20693 | 12001 | 99% | 57% |
| Equatorial Guinea | 27103 | 26715 | 99% | 98% |
| Eritrea | 121605 | 41211 | 75% | 25% |
| Estonia | 45792 | 38668 | 36% | 30% |
| Ethiopia | 1134770 | 837717 | 69% | 51% |
| Falkland Islands (Malvinas) | 11476 | 10703 | 62% | 57% |
| Faroe Islands | 1478 | 1101 | 33% | 25% |
| Fiji | 18132 | 14937 | 55% | 45% |
| Finland | 333890 | 301226 | 33% | 30% |
| France | 547871 | 363716 | 53% | 35% |
| French Guiana | 84152 | 83254 | 100% | 99% |
| French Polynesia | 2095 | 1395 | 0% | 0% |
| French Southern & Antarctic Lands | 7472 | 177 | 59% | 1% |

| Country | Area ($km^2$) | Total natural ecosystems ($km^2$) | Retention (% remaining natural ecosystems) | Retention (% land area) |
|---|---|---|---|---|
| Gabon | 262447 | 255104 | 99% | 96% |
| Gambia | 10786 | 4854 | 87% | 39% |
| Gaza Strip | 373 | 78 | 0% | 0% |
| Georgia | 70005 | 55853 | 90% | 72% |
| Germany | 356724 | 237181 | 56% | 37% |
| Ghana | 240579 | 171984 | 88% | 63% |
| Gibraltar | 8 | 4 | 72% | 34% |
| Greece | 130193 | 99936 | 78% | 60% |
| Grenada | 348 | 275 | 91% | 71% |
| Guadeloupe | 1661 | 1274 | 81% | 62% |
| Guatemala | 109632 | 90077 | 99% | 81% |
| Guernsey | 73 | 32 | 10% | 5% |
| Guinea | 246575 | 209721 | 91% | 78% |
| Guinea-Bissau | 33362 | 27236 | 69% | 57% |
| Guyana | 211982 | 202612 | 98% | 94% |
| Haiti | 27297 | 14457 | 97% | 51% |
| Honduras | 112855 | 97813 | 99% | 86% |
| Hungary | 92949 | 51476 | 43% | 24% |
| Iceland | 102510 | 69026 | 63% | 42% |
| India | 3166770 | 1120862 | 79% | 28% |

| Country | Area ($km^2$) | Total natural ecosystems ($km^2$) | Retention (% remaining natural ecosystems) | Retention (% land area) |
|---|---|---|---|---|
| Indonesia | 1905880 | 1608638 | 95% | 80% |
| Iran | 1627770 | 351847 | 41% | 9% |
| Iraq | 437470 | 80732 | 40% | 7% |
| Ireland | 69507 | 39991 | 28% | 16% |
| Isle of Man | 617 | 459 | 30% | 22% |
| Israel | 20771 | 4559 | 53% | 12% |
| Italy | 300155 | 215931 | 80% | 57% |
| Jamaica | 11092 | 9057 | 98% | 80% |
| Jan Mayen | 469 | 88 | 92% | 17% |
| Japan | 371862 | 305677 | 96% | 79% |
| Jersey | 124 | 29 | 7% | 2% |
| Jordan | 89489 | 8471 | 30% | 3% |
| Juan De Nova Island | 6 | 1 | 64% | 6% |
| Kazakhstan | 2721400 | 479518 | 30% | 5% |
| Kenya | 585741 | 389535 | 53% | 35% |
| Kiribati | 422 | 65 | 0% | 0% |
| Kuwait | 16797 | 329 | 15% | 0% |
| Kyrgyzstan | 199733 | 86095 | 75% | 32% |
| Laos | 231120 | 212735 | 99% | 91% |
| Latvia | 64469 | 55742 | 36% | 31% |

| Country | Area ($km^2$) | Total natural ecosystems ($km^2$) | Retention (% remaining natural ecosystems) | Retention (% land area) |
|---|---|---|---|---|
| Lebanon | 10240 | 7629 | 80% | 60% |
| Lesotho | 30621 | 16484 | 99% | 53% |
| Liberia | 96635 | 93720 | 98% | 95% |
| Libya | 1623940 | 17147 | 32% | 0% |
| Liechtenstein | 176 | 156 | 100% | 89% |
| Lithuania | 64858 | 48498 | 32% | 24% |
| Luxembourg | 2578 | 1927 | 67% | 50% |
| Macedonia | 25483 | 20159 | 83% | 66% |
| Madagascar | 596085 | 527377 | 92% | 81% |
| Malawi | 119199 | 55469 | 90% | 42% |
| Malaysia | 330691 | 311003 | 97% | 91% |
| Maldives | 34 | 1 | 9% | 0% |
| Mali | 1259190 | 293827 | 46% | 11% |
| Malta | 295 | 96 | 89% | 29% |
| Marshall Islands | 32 | 1 | 25% | 0% |
| Martinique | 1155 | 839 | 70% | 51% |
| Mauritania | 1043780 | 75239 | 30% | 2% |
| Mauritius | 2156 | 993 | 85% | 39% |
| Mayotte | 438 | 298 | 89% | 61% |
| Mexico | 1965660 | 1511500 | 75% | 58% |

| Country | Area ($km^2$) | Total natural ecosystems ($km^2$) | Retention (% remaining natural ecosystems) | Retention (% land area) |
|---|---|---|---|---|
| Midway Islands | 8 | 2 | 0% | 0% |
| Moldova | 33671 | 17356 | 82% | 42% |
| Monaco | 9 | 3 | 67% | 23% |
| Mongolia | 1562320 | 308055 | 59% | 12% |
| Montenegro | 13804 | 11997 | 93% | 81% |
| Montserrat | 113 | 61 | 89% | 48% |
| Morocco | 404446 | 133638 | 63% | 21% |
| Mozambique | 790638 | 709925 | 82% | 74% |
| Myanmar | 670372 | 533709 | 97% | 78% |
| Namibia | 827576 | 556207 | 46% | 31% |
| Nauru | 27 | 1 | 65% | 3% |
| Nepal | 147710 | 108751 | 99% | 73% |
| Netherlands | 35508 | 17884 | 31% | 16% |
| Netherlands Antilles | 794 | 259 | 66% | 22% |
| New Caledonia | 18938 | 17600 | 98% | 92% |
| New Zealand | 268943 | 192637 | 85% | 61% |
| Nicaragua | 128883 | 92155 | 99% | 70% |
| Niger | 1188520 | 55058 | 26% | 1% |
| Nigeria | 914291 | 461830 | 79% | 40% |
| Niue | 251 | 219 | 0% | 0% |

| **Country** | **Area ($km^2$)** | **Total natural ecosystems ($km^2$)** | **Retention (% remaining natural ecosystems)** | **Retention (% land area)** |
|---|---|---|---|---|
| Norfolk Island | 49 | 12 | 86% | 20% |
| North Korea | 122352 | 97303 | 95% | 76% |
| Norway | 319568 | 290047 | 73% | 66% |
| Oman | 310314 | 12987 | 71% | 3% |
| Pacific Islands (Palau) | 355 | 275 | 91% | 71% |
| Pakistan | 879493 | 205201 | 69% | 16% |
| Palestine | 5899 | 3665 | 57% | 35% |
| Panama | 74548 | 64819 | 98% | 85% |
| Papua New Guinea | 465554 | 450675 | 99% | 96% |
| Paraguay | 400654 | 324875 | 81% | 66% |
| Peru | 1299020 | 1055729 | 98% | 80% |
| Philippines | 294202 | 211471 | 95% | 69% |
| Poland | 311161 | 218737 | 54% | 38% |
| Portugal | 92027 | 75938 | 55% | 45% |
| Puerto Rico | 9197 | 7041 | 95% | 73% |
| Qatar | 11144 | 34 | 43% | 0% |
| Reunion | 2655 | 1804 | 93% | 63% |
| Romania | 237334 | 147574 | 68% | 43% |
| Russia | 16887500 | 15059036 | 85% | 76% |
| Rwanda | 25306 | 9790 | 94% | 36% |

| Country | Area (km²) | Total natural ecosystems (km²) | Retention (% remaining natural ecosystems) | Retention (% land area) |
|---|---|---|---|---|
| Saint Lucia | 639 | 571 | 84% | 75% |
| Samoa | 2955 | 2527 | 0% | 0% |
| San Marino | 60 | 33 | 92% | 51% |
| Sao Tome and Principe | 1150 | 848 | 97% | 72% |
| Saudi Arabia | 1963240 | 67606 | 28% | 1% |
| Senegal | 197173 | 135854 | 52% | 36% |
| Serbia | 88157 | 56556 | 74% | 47% |
| Seychelles | 380 | 139 | 91% | 33% |
| Sierra Leone | 72952 | 56061 | 95% | 73% |
| Singapore | 555 | 153 | 93% | 26% |
| Slovakia | 48885 | 35041 | 70% | 50% |
| Slovenia | 20416 | 18469 | 94% | 85% |
| Solomon Islands | 27157 | 24436 | 91% | 82% |
| Somalia | 640471 | 304232 | 33% | 16% |
| South Africa | 1225730 | 950442 | 55% | 43% |
| South Georgia and the South Sandwich Is | 3710 | 1597 | 0% | 0% |
| South Korea | 97428 | 77129 | 95% | 76% |
| Spain | 506729 | 374858 | 70% | 52% |
| Sri Lanka | 66468 | 51415 | 86% | 67% |

| Country | Area (km²) | Total natural ecosystems (km²) | Retention (% remaining natural ecosystems) | Retention (% land area) |
|---|---|---|---|---|
| St. Helena | 131 | 63 | 91% | 44% |
| St. Kitts and Nevis | 197 | 122 | 85% | 52% |
| St. Pierre and Miquelon | 223 | 173 | 94% | 73% |
| St. Vincent and the Grenadines | 344 | 317 | 89% | 82% |
| Sudan | 2501650 | 857130 | 48% | 16% |
| Suriname | 145952 | 143302 | 100% | 98% |
| Svalbard | 62048 | 30 | 84% | 0% |
| Swaziland | 17179 | 14182 | 94% | 78% |
| Sweden | 444324 | 405501 | 49% | 45% |
| Switzerland | 41472 | 31023 | 87% | 65% |
| Syrian Arab Republic | 188405 | 30246 | 25% | 4% |
| Taiwan | 36338 | 29950 | 98% | 81% |
| Tajikistan | 142644 | 64430 | 87% | 39% |
| Tanzania | 947556 | 676320 | 89% | 64% |
| Thailand | 515209 | 300730 | 91% | 53% |
| Togo | 57485 | 25222 | 92% | 41% |
| Tonga | 459 | 171 | 0% | 0% |
| Trinidad and Tobago | 5042 | 4259 | 92% | 78% |
| Tunisia | 155754 | 19155 | 23% | 3% |

| Country | Area ($km^2$) | Total natural ecosystems ($km^2$) | Retention (% remaining natural ecosystems) | Retention (% land area) |
|---|---|---|---|---|
| Turkey | 781034 | 496967 | 58% | 37% |
| Turkmenistan | 472324 | 122039 | 30% | 8% |
| Turks and Caicos Islands | 287 | 138 | 40% | 19% |
| Uganda | 243703 | 123937 | 80% | 41% |
| Ukraine | 596959 | 311595 | 25% | 13% |
| United Arab Emirates | 70631 | 1899 | 37% | 1% |
| United Kingdom | 243845 | 131799 | 55% | 30% |
| United States | 9470940 | 6082574 | 73% | 47% |
| Uruguay | 178357 | 44695 | 82% | 20% |
| Uzbekistan | 446967 | 93475 | 30% | 6% |
| Vanuatu | 12335 | 10561 | 77% | 66% |
| Venezuela | 916780 | 832227 | 93% | 85% |
| Vietnam | 326038 | 193177 | 99% | 59% |
| Virgin Islands | 219 | 106 | 43% | 21% |
| Wake Island | 18 | 3 | 38% | 5% |
| Wallis and Futuna | 161 | 5 | 0% | 0% |
| Western Sahara | 270160 | 457 | 20% | 0% |
| Yemen | 426123 | 69984 | 70% | 12% |
| Zambia | 756427 | 693771 | 89% | 81% |

| **Country** | **Area (km$^2$)** | **Total natural ecosystems (km$^2$)** | **Retention (% remaining natural ecosystems)** | **Retention (% land area)** |
|---|---|---|---|---|
| Zimbabwe | 391930 | 340713 | 80% | 69% |
